# Not all TOP RNAs are created equal: 3’UTR length and TSS selection predict the translational regulation of LARP1-bound mRNAs in CD4^+^ T cells

**DOI:** 10.64898/2026.09.25.754325

**Authors:** Alison Galloway, Victoria H. Cowling

**Affiliations:** Cancer Research UK Scotland Institute, Switchback Road, Glasgow, G61 1BD, UK; School of Life Sciences, University of Dundee, Dundee, DD1 5EH, UK; School of Cancer Sciences, University of Glasgow, Switchback Road, Glasgow, G61 1QH, UK

## Abstract

Naïve T cells are poised for activation and contain a pool of translationally repressed ribosomal protein (RP) mRNA prepared to induce ribosome biogenesis to support protein synthesis, cell growth and proliferation. RP mRNA are the prototypical members of a class of transcripts initiating at cytosine followed by a CU rich element called terminal oligo pyrimidine (TOP) RNAs. TOP RNAs are regulated by an RNA binding protein LARP1, which promotes transcript stabilisation and translational repression. We investigated LARP1 function in T cell activation by generating cross-linking immunoprecipitation (CLIP) datasets detailing the LARP1-RNA interactions in naïve and activated CD4^+^ T cells and identifying novel TOP RNAs. TOP RNAs identified by this analysis were functionally diverse. RP mRNAs were typified by high stability, and translational repression in naïve T cells followed by MTORC1-dependent translation increases following T cell activation. However, other TOP RNAs varied in these aspects of their regulation. Notably, TOP RNAs with longer 3’UTRs had a relaxed dependency on LARP1 for stability and a reduced dependency on MTORC1 for their translation. Transcription start site heterogeneity also impacted TOP RNA regulation by generating a mixture of transcript isoforms with different TOP motif lengths. Longer terminal oligo pyrimidine stretches were associated with a greater dependency on MTORC1 for translation. Differential regulation of TOP RNAs may allow tuneable translational responses to MTORC1 and indicates potential roles for LARP1 beyond translation regulation and stability.

## Introduction

To allow the detection of novel pathogens and cancer neoantigens the immune system generates a large pool of naïve T cells with differing T cell receptor (TCR) specificities. These naïve T cells remain quiescent until activation is triggered by stimulation of the TCR by an antigen presenting cell. Following activation, T cells rapidly increase their biosynthetic capacity, resulting in increased cell size and rapid proliferative expansion (Howden et al. 2019; Chapman et al. 2020). They then differentiate into effector T cells that perform immune functions, including direct killing and the secretion of cytokines that signal to other immune cells. To allow this rapid transition from the quiescent state, naïve T cells remain poised for activation with certain mRNAs, including ribosomal protein (RP) mRNAs, being pre-synthesised, but translationally repressed (Tan et al. 2017b; Wolf et al. 2020; Turner 2023).

mRNAs encoding RPs are very abundant in naive T cells making up approximately a quarter of the protein-coding transcriptome(Galloway et al. 2021). These RP transcripts are translationally repressed in naïve T cells, then following TCR stimulation, mammalian target of rapamycin complex 1 (MTORC1) signalling promotes their translation (Tan et al. 2017a; Wolf et al. 2020). This mechanism allows a rapid induction of ribosome biogenesis following T cell activation. RP transcripts belong to a class of RNAs that initiate with a cytosine followed by ∼10 pyrimidines called terminal oligopyrimidine (TOP) RNAs (Meyuhas and Kahan 2015). The TOP motif interacts with the RNA binding protein La-related protein 1 (LARP1) which stabilises TOP RNAs and regulates their translation (Fonseca et al. 2018; Berman et al. 2021). LARP1 has multiple RNA binding domains, the DM15 domain interacts with the RNA cap and first four nucleotides (Lahr et al. 2015; Lahr et al. 2017; Cassidy et al. 2019), the La-module interacts with both the TOP motif and the polyA tail (Aoki et al. 2013; Al-Ashtal et al. 2021), and the ribosome binding domain interacts with the 40S ribosomal subunit (Saba et al. 2024; Wolin et al. 2025). When MTORC1 is inactive, LARP1 represses the translation of RP mRNAs, then under conditions of cell growth, phosphorylation of LARP1 by MTORC1 causes LARP1 to release the RNA cap to promote the translation of RPs (Tcherkezian et al. 2014; Fonseca et al. 2015; Hong et al. 2017; Jia et al. 2021). LARP1-dependent maintenance of a stabilised, but translationally repressed pool of RP transcripts under conditions of MTORC1 repression allows a more rapid recovery of ribosome biogenesis and protein translation capacity when MTORC1 signalling is restored (Fuentes et al. 2021). Consistent with an important role for the RNA cap-LARP1 interaction in RP mRNA stabilisation, they also have enhanced dependence on the RNA cap methyltransferase enzyme RNMT and its cofactor RAM (RNMT activating miniprotein) for their expression. RNMT and RAM are induced following T cell activation and promote ribosome biogenesis and proliferation in activated T cells(Galloway et al. 2021; Knop et al. 2024).

The rapid induction of ribosome biogenesis following T cell activation thus provides a physiologically important platform to investigate TOP RNA regulation (Asmal et al. 2003; Allison et al. 2016; Tan et al. 2017b; Galloway et al. 2021; Rosenlehner et al. 2024). Here we investigate LARP1 expression, ribosome association and RNA binding in CD4^+^ T cells. We use cross linking immunoprecipitation (CLIP) datasets to map the LARP1-RNA interactions in naïve and activated CD4^+^ T cells, identifying novel TOP RNAs including those with lower abundance or T cell specific expression including the TCR components Cd247 (Cd3ζ), Cd3δ and Cd4. We then investigate the regulation of these functionally diverse TOP RNAs across T cell activation. We find that RP transcripts undergo typical TOP RNA regulation, with high transcript stability, and translational repression in naïve T cells that is lifted in an MTORC1-dependent manner following T cell activation. In contrast, many other TOP RNAs adhere to these rules less stringently. RP transcripts have very short 3’UTRs and more stringent TOP motifs, both of these factors have been linked to their translational regulation in experiments using reporter RNAs (Ledda et al. 2005; Philippe et al. 2020). Here we find that short 3’UTR length and higher quality TOP motifs correlate with greater translational repression of TOP mRNAs in naïve T cells, greater MTORC1-dependent induction of protein expression upon activation and a greater effect of LARP1 on transcript stability. Thus, we have uncovered a diversity in TOP RNA function and regulation in T cells.

## Results

### LARP1 is induced during T cell activation and is associated with ribosomes

T cell activation, triggered by T cell receptor (TCR) ligation, induces cell growth, proliferation and differentiation into effector T cells. To examine the role of LARP1 in T cell activation we first investigated the expression and localisation of LARP1 in mouse CD4^+^ T cells. Larp1 mRNA and protein expression was substantially induced during CD4^+^ T cell activation, with the protein increasing approximately threefold within 16 hours (Fig.1A-C) (Tan et al. 2017a). This change in expression occurs alongside increases in transcription and ribosome biogenesis, thus correlates with an increased need to upregulate TOP RNAs.

**Figure 1:**
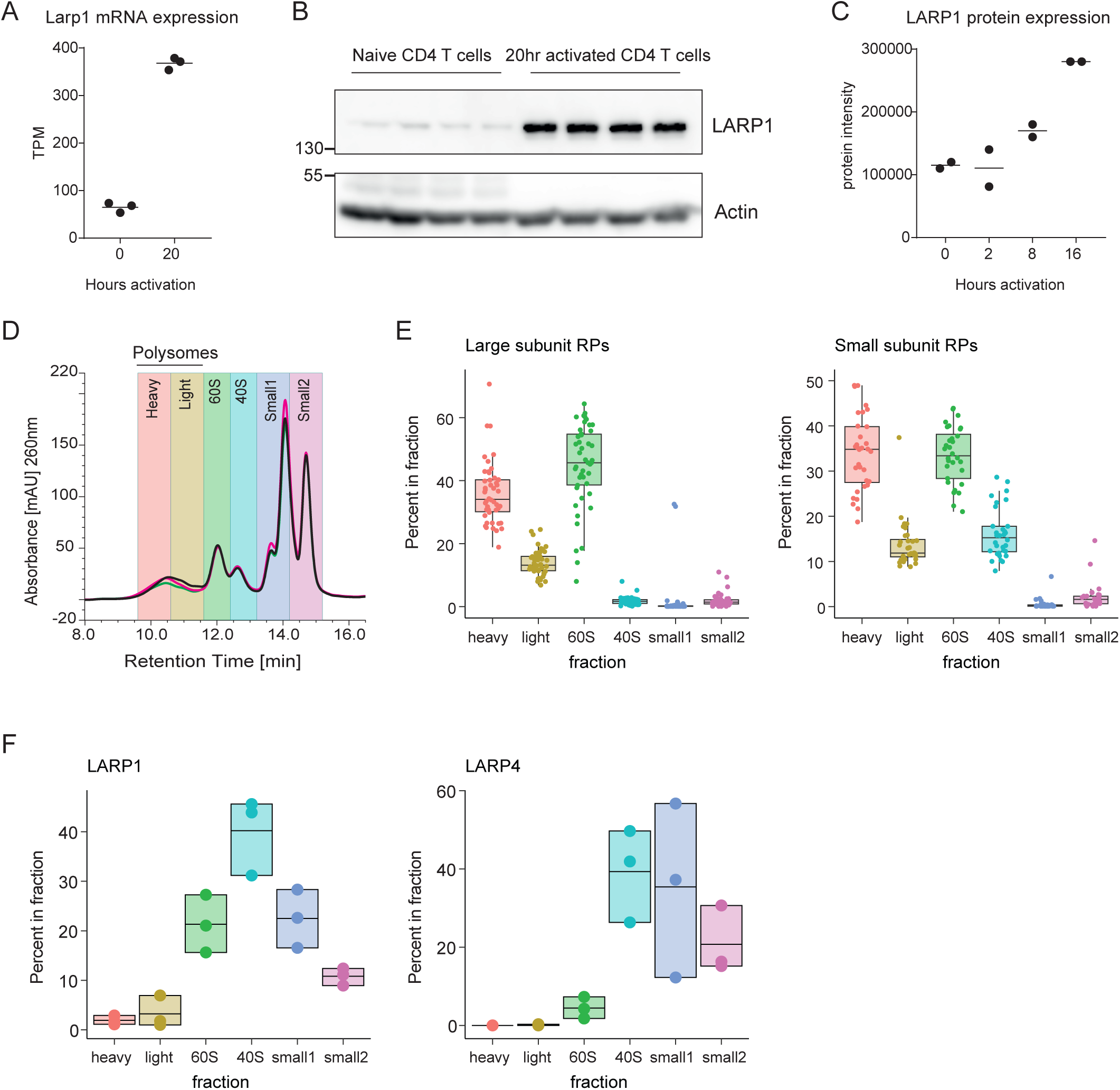
LARP1 associates with ribosomes in activated CD4^+^ T cells. **(A)** Transcripts per million (TPM) quantitation of Larp1 mRNA form RNAseq data of naïve (n=3 mice) and 20h activated (n=3 mice) CD4^+^ T cells. Each point represents a biological replicate, line represents the mean. **(B)** Western blot showing expression of LARP1 and Actin in naïve (n=4 mice) and 20 hour activated (n=4 mice) CD4^+^ T cells. 1 million cells were loaded per well, both proteins detected on same membrane. **(C)** Proteomics analysis of LARP1 expression in naïve and activated mouse CD4^+^ T cells(Tan et al. 2017a) each point represents a biological replicate (n= 2 mouse samples). **(D)** Polysomes and ribosomes from 20 hour activated CD4 T cell lysates (n=3 mice) were resolved by Ribo mega-SEC; 260nm UV absorbance profile showing the fractions collected. **(E)** Fractions from the Ribo Mega-SEC were analysed by label free proteomics. iBAQ intensities for each protein were normalised to the total iBAQ protein intensity from that fraction. Mean normalised intensities within each fraction are plotted for RPS and RPL proteins. Each point represents a protein, boxes show 25^th^, 50^th^ and 75^th^ percentile, whiskers represent the largest/smallest values no further than 1.5 X the interquartile range from the box. **(F)** Analysis of the distribution of LARP family members in the CD4 T cell Ribo Mega-SEC fractions. Points represent the normalised protein intensities for each biological replicate (n=3 mice), and boxes show the range and mean normalised protein intensity values.

Prior research indicates that LARP1 interacts with the 40S ribosomal subunit (Gentilella et al. 2017; Saba et al. 2024; Wolin et al. 2025). We resolved activated CD4^+^ T cell polysomes and ribosomes by Ribo Mega-SEC, a technique for the separation of large complexes by size exclusion chromatography (Fig.1D) (Yoshikawa et al. 2018; Yoshikawa et al. 2021). Proteins from each fraction were analysed by mass spectrometry (Table S1). Small and large subunit RPs indicated were enriched in the polysomal and 40S and 60S ribosomal fractions (Fig.1E). The 60S fraction contained large and small subunit RPs, indicating overspill of the 40S or 80S ribosomal subunits into this fraction, whereas the 40S fraction was only enriched for small subunit RPs. Two proteins in the La-related protein family, LARP1 and LARP4, were detected in the Ribo Mega-SEC fractions. LARP1 and LARP4 are known to interact with the 40S ribosomal subunit (Yang et al. 2011; Gentilella et al. 2017; Saba et al. 2024; Ranjan et al. 2025; Wolin et al. 2025) and were enriched in the 40S fraction in activated CD4^+^ T cells (Fig.1F). LARP1 was also detected, at lower levels, in the polysomal fractions and in the smaller protein complex fractions where it could be in complexes with RNA, proteins or unbound. Thus, in activated CD4^+^ T cells, LARP1 expression is induced and a large proportion of the LARP1 protein co-localises with the 40S ribosomal subunit.

### LARP1-RNA interaction analysis (CLIP) identifies functionally diverse TOP RNAs

LARP1 has multiple RNA binding domains and their association with RNA may be regulated during T cell activation by post transcriptional modifications or by competition with other RNA binding proteins (Kakegawa et al. 2007; Damgaard and Lykke-Andersen 2011). We assessed direct LARP1-RNA interactions in naïve and activated CD4^+^ T cells by cross linking immunoprecipitation (CLIP), a method that identifies the direct binding of proteins to RNA with nucleotide level precision (Van Nostrand et al. 2017; Galloway et al. 2021). Consistent with LARP1 binding to the TOP motif, the majority of LARP1 binding sites, identified using DEWseq (Sahadevan et al. 2022; Schwarzl et al. 2024), were in the 5’UTRs of protein coding transcripts (Fig.2A, Table S2) and the most common motif was CU rich (Fig.2B, Table S3). There were also LARP1 binding sites located in long non-coding RNAs (lncRNA) of which 13 were sno-RNA host transcripts (Fig.2A, Table S2). The majority of RNA-binding occurred adjacent to the transcript start site implying the RNA binding observed may involve the cap-dependent DM15 domain (Fig.2C-D). LARP1 also interacted with a binding site on the 18S rRNA around nucleotide 1703 (Fig.2E) confirming a direct interaction with the 40S ribosomal subunit in T cells (Saba et al. 2024; Wolin et al. 2025).

**Figure 2.**
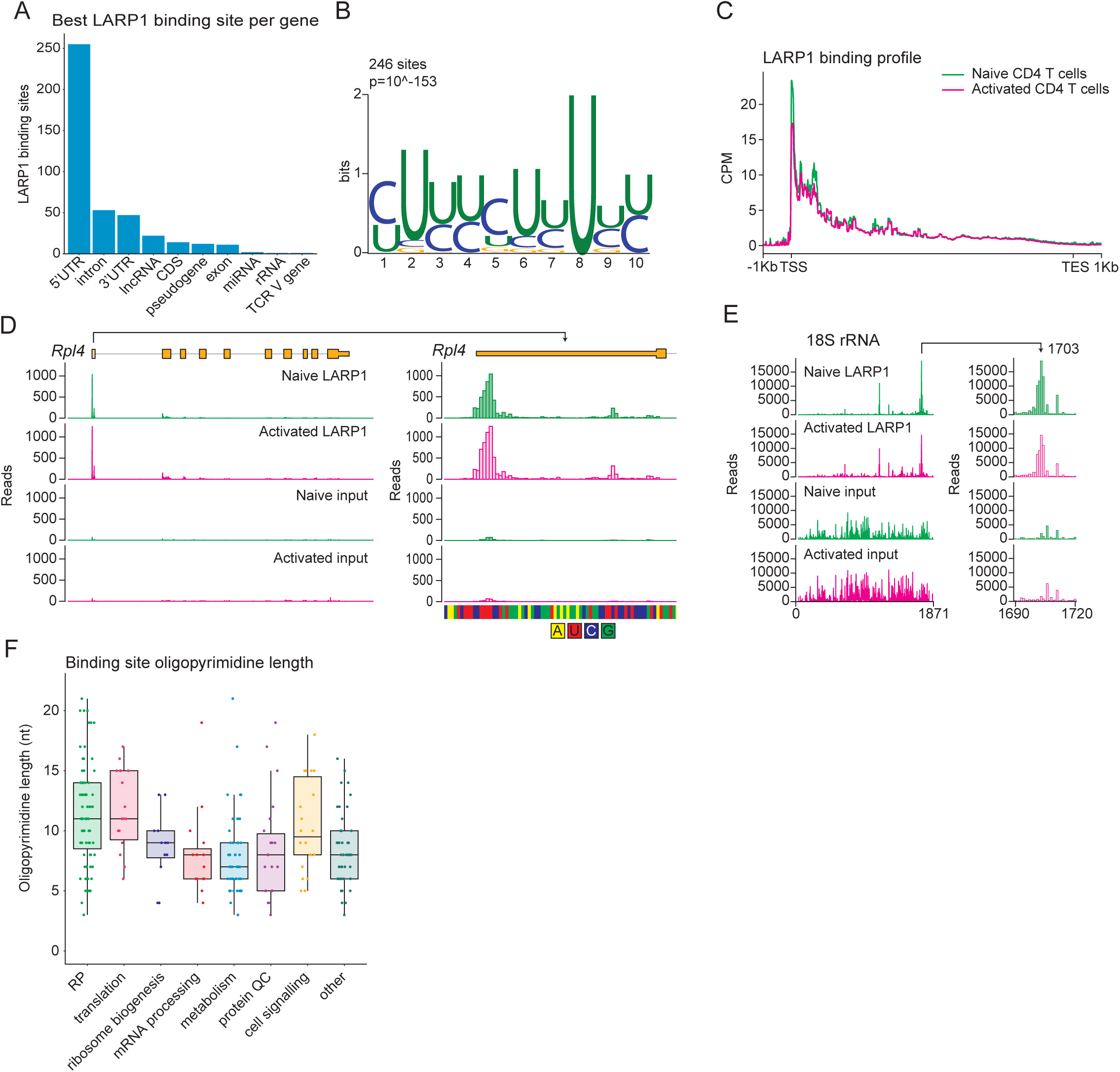
LARP1 directly binds to the TOP motifs at the 5’ end of transcripts. **(A-B)** LARP1 binding sites were determined in two mouse naïve CD4^+^ T cell datasets using DEWseq, which uses a sliding window approach to compare LARP1 CLIP and input data. For each transcript with a binding site, the window with the lowest p-value was selected, the transcript features that are overlapped by these best windows are quantified in (A). Sequences from the best windows were analysed by MEME to generate the most common motif (B). **(C)** Distribution of naïve and activated CD4^+^ T cell LARP1 CLIP reads across transcripts where LARP1 binding sites were identified. Transcripts were length normalised between the transcript start site (TSS) and transcript end site (TES). CPM= counts per million mapped. **(D-E)** Distribution of naïve and activated CD4^+^ T cell LARP1 CLIP reads across the *Rpl4* gene (D) and 18S rRNA (E). Reads indicate the number of reads starting at that position; read starts are expected to be 1nt downstream of the crosslink site, yellow shapes represent the exons, lines represent the introns. **(F)** Longest oligopyrimidine stretch within the best window (lowest p-value from DEWseq analysis) for each protein-coding TOP RNA defined as the transcripts with LARP1 binding sites within the 5’UTR. TOP RNAs were grouped according to function. Each point represents one gene, boxes show 25^th^, 50^th^ and 75^th^ percentile, whiskers represent the largest/smallest values or 1.5 X the interquartile range.

Since they made up the majority of hits, we focussed our analysis on TOP RNAs, which we defined as transcripts with a LARP1 binding region identified within the 5’UTR (Table S4). While TOP RNAs can be defined directly based on their sequences, which begin with a string of pyrimidines, using the LARP1 binding data circumvents challenges with transcription start site heterogeneity and instead relies on the interaction between LARP1 and TOP sequences (Fig.2B-D). TOP RNAs defined by LARP1 binding typically had a pyrimidine stretch between 5 and 15 nucleotides long within the most significant LARP1 binding region (Fig.2F). We used STRING analysis (Szklarczyk et al. 2023) to illustrate the functional diversity of the 265 TOP RNAs identified (Fig. 3A). In addition to the ribosome, we found that TOP RNAs encode components of other multi-protein complexes such as the electron transport chain complexes I (NADH dehydrogenase), III (ubiquinone-cytochrome c reductase), IV (cytochrome c oxidase) and V (ATP synthase). Outside of these larger complexes, many proteins encoded by TOP RNAs are functionally related. For example, TOP RNAs encode the outer mitochondrial membrane transporters TOMM20 and TOMM7, selected mitochondrial ribosomal proteins, mitochondrial solute carries SLC25A3 (Solute Carrier Family 25 Member 3) and SLC25A30 (Solute Carrier Family 25 Member 30) and cristae junction proteins MICOS13 (Mitochondrial Contact Site And Cristae Organizing System Subunit 13) and CHCHD3 (Coiled-Coil-Helix-Coiled-Coil-Helix Domain Containing 3), which could be co-ordinately regulated with the electron transport chain components. However, there are also many TOPs encoding proteins with no direct relations.

**Figure 3.**
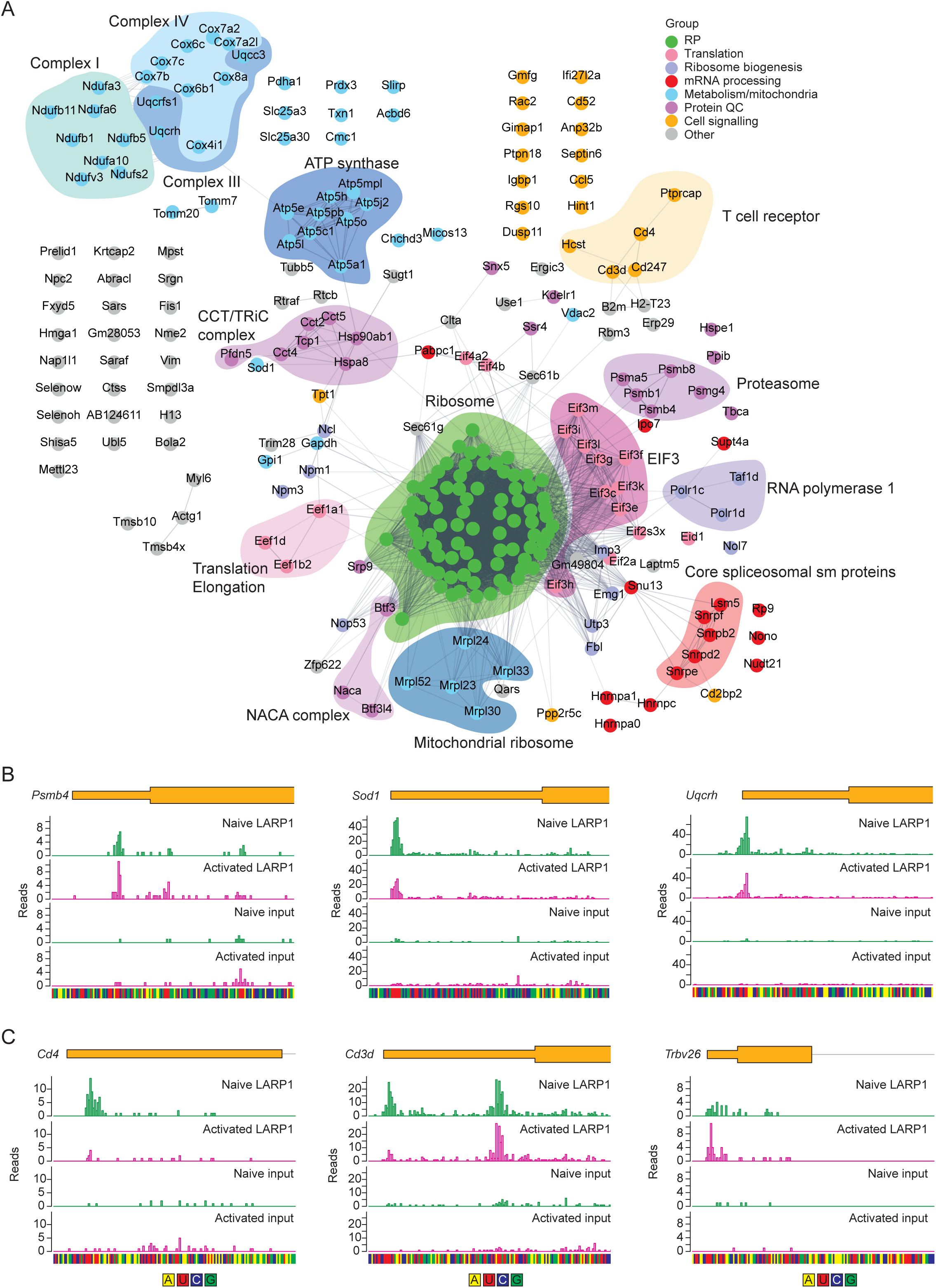
LARP1 binds to functionally diverse TOP RNAs. **(A)** STRING analysis of TOP RNAs identified by LARP1 CLIP analysis. Each node represents a protein encoded by a TOP RNA, each line indicates a physical interaction between proteins. **(B-C)** Distribution of naïve and activated CD4^+^ T cell LARP1 CLIP reads across non-RP (B) and T cell specific (C) transcripts. Reads (y-axis) indicate the number of reads starting at that position; read starts are expected to be 1nt downstream of the crosslink site, yellow shapes represent the exons, lines represent the introns.

CLIP data demonstrating LARP1 binding to example non-RP TOP RNAs is shown in figure 3B-C, and as with other TOPs (Fig.2C-D), occurs at the 5’ end of the transcript to CU rich sequences. Interestingly, we identified a group of T cell-specific TOP transcripts comprised of TCR components including: Cd247 (Cd3ζ), Cd3d (Cd3δ), Cd4 and, surprisingly, one of the TCRβ variable region segments (Trbv26). There were changes in the amount and exact position of LARP1 binding detected on these transcripts between naive and activated CD4^+^ T cells suggesting there may be dynamic regulation of LARP1 binding to these transcripts across T cell activation (Fig.3C).

### LARP1-RNA interactions are maintained following T cell activation

Previous studies indicate that in growth promoting conditions, LARP1 releases the RNA cap allowing EIF4E to bind, increasing the translation of TOP RNAs (Tcherkezian et al. 2014; Fonseca et al. 2015; Hong et al. 2017; Jia et al. 2021). To determine whether the distribution of LARP1 between different TOP transcripts changed following activation we compared total RNAseq and LARP1 CLIP read densities in naive and activated CD4^+^ T cells (Fig.4A-B). Our CLIP analysis used a sliding windows approach, so for each gene the window with the lowest P value for the comparison between LARP1 CLIP and the size matched input was selected to represent that gene. The density of LARP1 CLIP reads was proportional to the overall abundance of each target transcript (Fig.4A-B). However, the ratio of CLIP reads to total transcripts per million is highest for RPs, particularly in activated CD4^+^ T cells, suggesting a larger proportion of these transcripts are interacting with LARP1 (Fig.4C-D). This broad analysis does not preclude subtle changes: notably activation drives a major remodelling of the CD4^+^ T cell transcriptome including a three-fold increase in mRNA per cell making normalisation non-trivial (Galloway et al. 2025). In accordance with their high expression level, we estimate from this data that more than 70% of LARP1-RNA interactions are with RP transcripts, this is important to consider when interpreting data related to LARP1 protein interactions and localisation.

**Figure 4.**
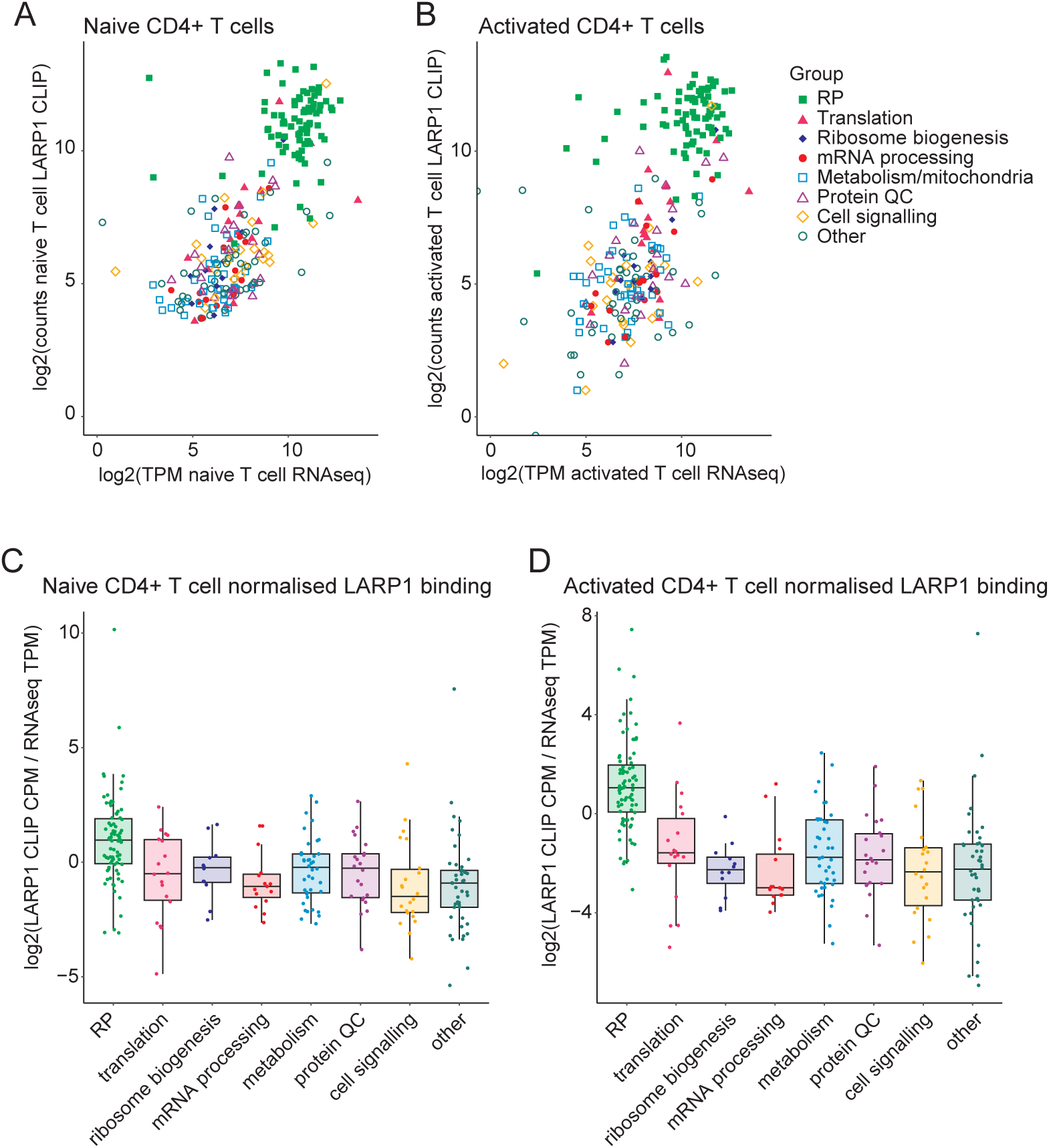
LARP1-RNA interactions are maintained following T cell activation. **(A-B)** Naïve (A) and activated (B) CD4^+^ T cell TOP RNA LARP1 CLIP reads starting within the best windows were quantified and normalised to total library size and then compared to total RNAseq (transcripts per million:TPM). Selected transcripts were assigned a function and coloured as shown. Each point represents one gene. **(C-D)** Naïve (C) and activated (D) CD4^+^ T cell TOP RNA LARP1 CLIP reads starting within the best windows were quantified and normalised to total library size and then to transcripts per million in the total RNAseq. Each point represents one gene, boxes show 25^th^, 50^th^ and 75^th^ percentile, whiskers represent the largest/smallest values or 1.5 X the interquartile range.

### TOP RNAs with shorter 3’UTRs have lower translation efficiency

RP transcripts are translationally repressed in naïve T cells, and their translation is induced following T cell activation to promote ribosome biogenesis(Wolf et al. 2020). However, in addition to RPs we identified TOP RNAs with a variety of cellular functions whose regulation is unknown. To determine how different TOP RNAs are regulated in T cells we divided them into functional groups and examined their translation efficiency based on ribosome footprinting analysis. We defined translational efficiency as the CPM of the ribosome protected fragments (RPFs) divided by the CPM for the total RNA for each gene. Within every functional group, the TOP mRNAs tended to be translationally repressed in naïve T cells, however, TOP RNAs encoding RPs had the strongest translational repression (Fig.5A) (Khalsa et al. 2019).

**Figure 5.**
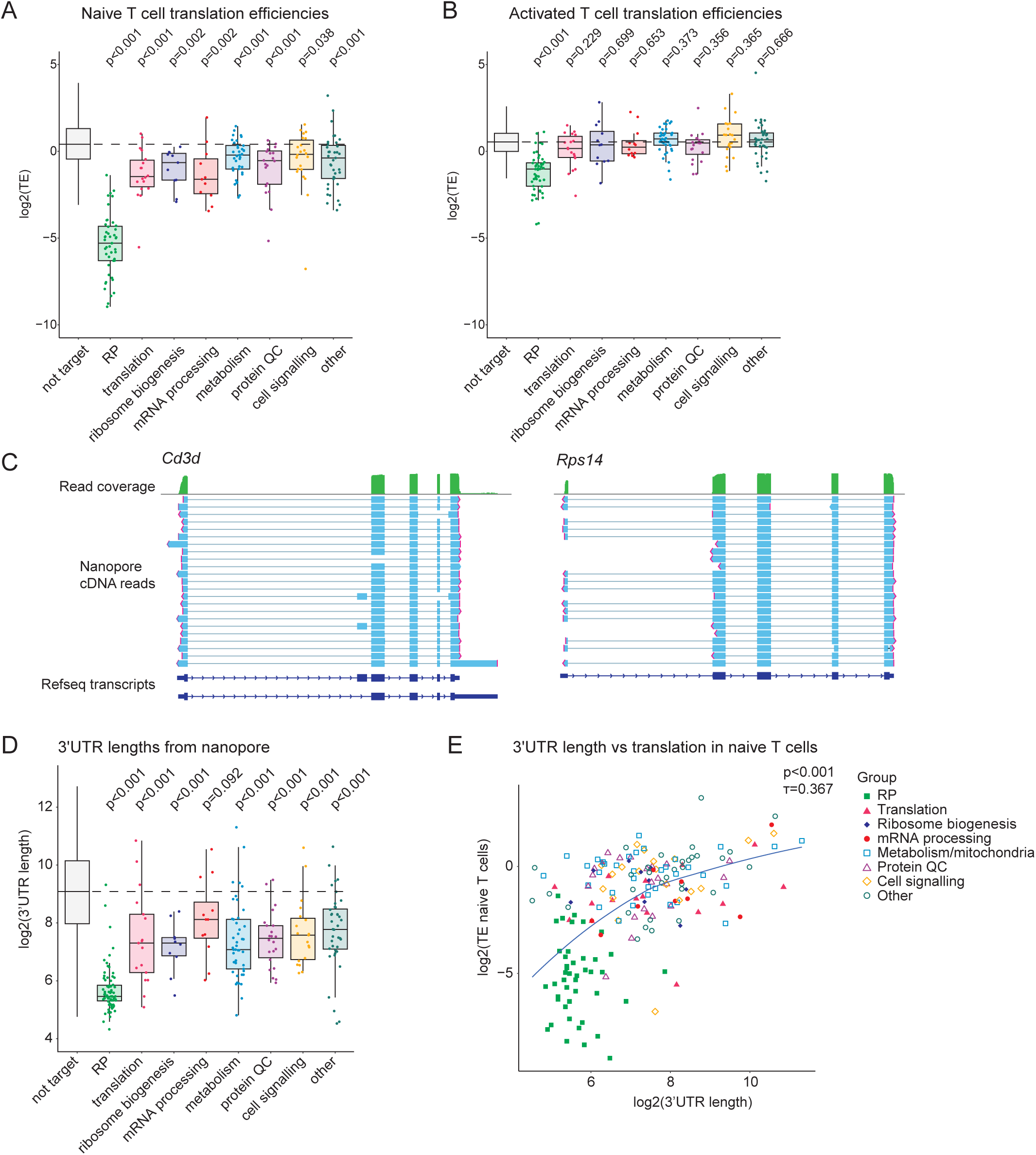
TOP RNA translation efficiency is associated with 3’UTR length. Protein-coding transcripts with LARP1 binding sites in the 5’UTR were divided into groups depending on their function. Transcripts lacking these LARP1 binding sites (not target) are shown for comparison. **(A)** Mouse naïve T cell translation efficiencies of LARP1-bound RNAs (average of 2 biological replicates)(Khalsa et al. 2019). **(B)** Mouse activated CD4^+^ T cell translation efficiencies of LARP1-bound RNAs (average of 3 biological replicates)(Galloway et al. 2021). **(C)** Full length cDNA Nanopore reads from mouse 20 hour activated CD4^+^ T cells, aligning to *Cd3d* and *Rps14* genes(Galloway et al. 2025). **(D)** 3’UTR length of LARP1-bound RNAs determined from the highest abundance transcript isoform of each gene in 20 hour activated CD4^+^ T cell full length cDNA Nanopore sequencing data. **(E)** Comparison of 3’UTR length with naïve T cell translation efficiency. TE (translation efficiency: ribosome protected fragment RNA counts per million (CPM)/total mRNA CPM). (A,B,D,E) Each point represents a gene, (A,B,D) boxes show 25^th^, 50^th^ and 75^th^ percentile, whiskers represent the largest/smallest values or 1.5 X the interquartile range. (A,B,D) TOP RNAs compared to “not target” group using Wilcoxon rank sum test with Benjamini-Hochberg p- value adjustment for multiple testing. (E) Correlation assessed by Kendall’s rank τ coefficient, trendline plotted using locally estimated scatterplot smoothing.

Following T cell activation, the translation efficiency of TOP RNAs tended to increase and only the RP transcripts still had low translation efficiency relative to non-TOP transcripts (Fig.5B). The low translation efficiency of RP transcripts in activated CD4^+^ T cells was surprising since these cells are undergoing a substantial increase in ribosome biogenesis. However, as well as increased ribosome biogenesis, the cells also undergo increased mRNA transcription, therefore, translational regulation of the increased pool of RP mRNA may remain important to avoid synthesising more RPs than required. This data is also consistent with the finding that the presence of LARP1 leads to translational repression of RP transcripts even in conditions of full MTOR activation (Jia et al. 2021).

The translational efficiency of non-RP TOP RNAs was significantly higher than that of RPs in both naïve and activated T cells. Whilst multiple factors can influence translation efficiency, we were intrigued by a study by Ledda *et al* demonstrating that translational repression of a RPS6 5’UTR reporter mRNA in serum starved cells depended on the length of the 3’UTR (Ledda et al. 2005). Alternative polyadenylation and cleavage sites can result in multiple transcript isoforms with different 3’ UTR lengths. Therefore, we used Nanopore direct cDNA sequencing data from activated CD4^+^ T cells (Galloway et al. 2025) to investigate 3’UTR length. Notably certain TOP transcripts had multiple transcript isoforms in T cells as shown for Cd3d (Fig.5C), whereas others such as Rps14 had only one isoform detected (Fig.5C). We determined the most commonly expressed 3’UTR for each TOP transcript within T cells (Table S5). Whilst TOP RNAs within most functional groups had shorter than average 3’UTRs, RPs had the shortest with a median length of only 44 nucleotides compared with 538 nucleotides for non-TOP transcripts (Fig.5D). Shorter TOP RNA 3’UTR length correlated with lower translational efficiency in naive T cells (Fig.5E). RP mRNAs had both the lowest translational efficiency and shortest 3’UTRs, therefore, we also assessed the relationship between TOP mRNA translation and 3’UTR length with RP mRNAs excluded and found that the correlation was still significant (Fig.S1A).

### TSS heterogeneity impacts TOP sequence quality and translation efficiency

An additional factor that can influence translational repression of TOP RNAs is transcript start site (TSS) heterogeneity resulting in different sequences at the 5’ end of the RNA. Whilst all TOP RNAs identified by LARP1 CLIP had oligo pyrimidine stretches within the LARP1 binding site (Fig.2F), only transcripts beginning in a pyrimidine are classed as TOP transcripts. Therefore, transcripts derived from the same gene can include a mixture of TOP and non-TOP isoforms and may have TOP motifs of different lengths depending on the exact site of transcription initiation. TSSs can be determined using cap analysis of gene expression (CAGE). The TOP score metric, developed by Phillipe *et al*, is the mean length of the consecutive pyrimidines at the start of a gene’s transcripts (Philippe et al. 2020). We used naïve mouse CD4^+^ T cell CAGE data from the FANTOM5 project to calculate TOP scores for each expressed gene (Fig.6A, Table S6) (Noguchi et al. 2017). TOP RNAs identified by LARP1 CLIP had significantly higher TOP scores than non-TOP RNAs. Within the TOP transcripts, those encoding RPs and translation factors had the highest scores. Consistent with the requirement of TOP sequences for LARP1 binding, we found that the TOP score correlated with the ratio of CLIP reads to total transcripts per million: an approximation of LARP1 binding efficiency (Fig.6B). Transcripts with higher TOP scores have been shown to have higher translational repression in Torin1 treated HEK 293T cells (Philippe et al. 2020). Here we found that higher TOP scores correlated with lower translation efficiency in the naïve T cells (Fig.6C, Fig.S1B).

**Figure 6.**
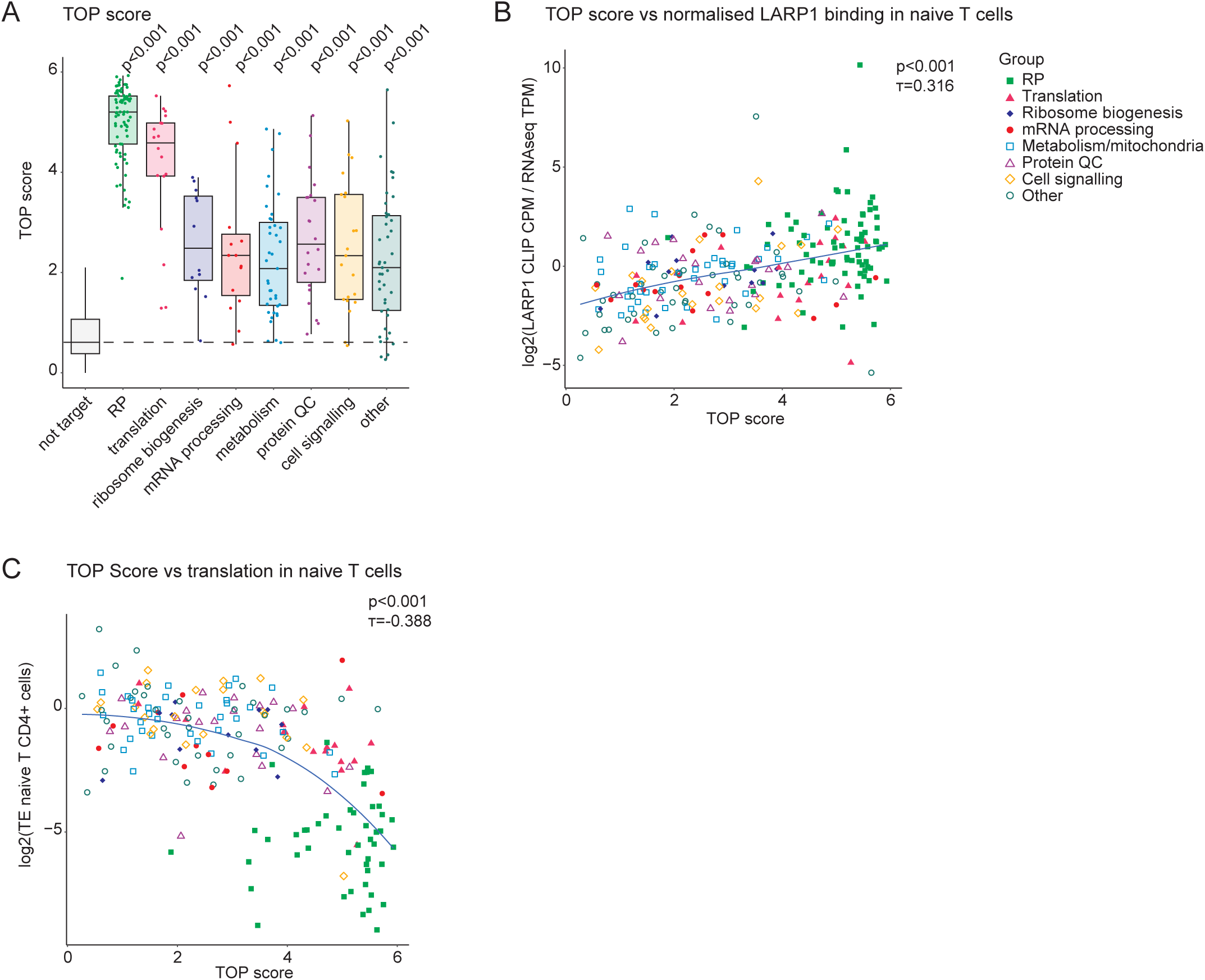
TOP score is associated with LARP1 binding and RNA translation efficiency. Protein-coding transcripts with LARP1 binding sites in the 5’UTR were divided into groups depending on their function. Transcripts lacking these LARP1 binding sites (not target) are shown for comparison. **(A)** TOP score (average 5’ terminal pyrimidine length) of LARP1-bound RNAs determined from FANTOM5 naïve CD4^+^ T cell CAGE data(Noguchi et al. 2017). Boxes show 25^th^, 50^th^ and 75^th^ percentile, whiskers represent the largest/smallest values or 1.5 X the interquartile range. TOP RNAs compared to “not target” group using Wilcoxon rank sum test with Benjamini-Hochberg p-value adjustment for multiple testing. **(B)** Comparison of TOP score with naïve CD4^+^ T cell LARP1 binding, determined by LARP1 CLIP normalised to total RNA. **(C)** Comparison of TOP score with naïve T cell translation efficiency. TE (translation efficiency: ribosome protected fragment RNA counts per million (CPM)/total mRNA CPM). (A,B,C) Each point represents a gene. (B,C) Correlation assessed by Kendall’s rank τ coefficient, trendline plotted using locally estimated scatterplot smoothing.

### TOP RNAs with shorter 3’UTRs are more dependent on LARP1 for stabilisation

A major role of LARP1 is to stabilise TOP transcripts, therefore we assessed the RNA half-lives of TOP RNAs in naive T cells using data from Hwang *et al*(Hwang et al. 2020). As expected, RP mRNAs were very stable (Fig.7A). Other TOP RNAs, however, varied in their stability, and this was related to transcript function. For instance, mRNAs encoding proteins involved in cell signalling typically have shorter half-lives to allow cells to rapidly adapt to changing conditions (Raghavan et al. 2002) and this was true of the TOP RNAs involved in cell signalling. The impact of LARP1 on transcript stability was measured in HeLa cells using SLAMseq (4-thiouracil pulse-chase labelling) by Hochstoeger *et al* (Hochstoeger et al. 2024). LARP1 increased the stability of TOP RNAs within each functional category, however, it had the greatest impact on RP transcripts (Fig.7B). We next investigated whether 3’UTR length was associated with TOP RNA stability. TOP transcripts with the shortest 3’UTRs had the greatest decrease in RNA stability in *LARP1* KO HeLa cells, whereas those with longer 3’UTRs were less affected (Fig.7C, Fig.S1C). TOP transcripts with higher TOP scores also had greater decreases in RNA stability in *LARP1* KO HeLa cells (Fig.7D). However, this was primarily driven by RP transcripts implying that the effect may be indirect (Fig.S1D). Since RPs had the shortest 3’UTRs as well as the highest TOP scores, this could explain the relationship between TOP score and stability. Consistent with this, TOP scores inversely correlated with 3’UTR length (Fig.7E), but only when RPs were included (Fig.S1E). Therefore, 3’UTR length correlates with TOP stabilisation by LARP1, whereas both 3’UTR length and TOP score correlate with low translational efficiency.

**Figure 7.**
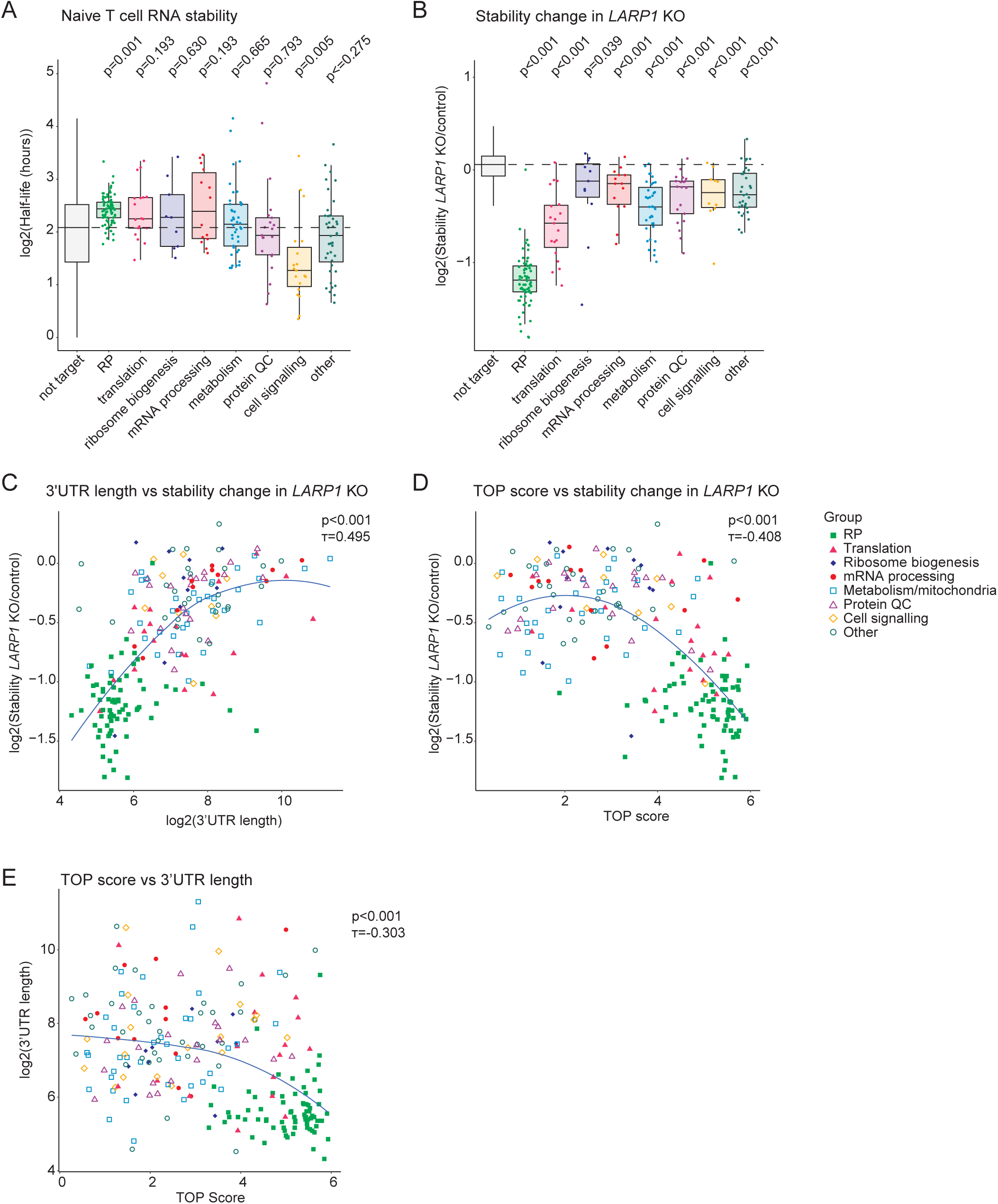
TOP RNA stabilisation is associated with shorter 3’UTRs. Protein-coding transcripts with LARP1 binding sites in the 5’UTR were divided into groups depending on their function. Transcripts lacking these LARP1 binding sites (not target) are shown for comparison. **(A)** Transcript stabilities of LARP1-bound RNAs in naïve CD4^+^ T cells(Hwang et al. 2020). **(B)** Change in transcript stability in *LARP1* KO HeLa cells (average of n=2 independent clones)(Hochstoeger et al. 2024). **^(C)^** Comparison of 3’UTR length determined from Nanopore sequencing data in activated CD4^+^ T cells with change in stability in *LARP1* KO HeLa cells. **(D)** Comparison of TOP score (average terminal pyrimidine length) determined from FANTOM5 naïve CD4 cell CAGE data with change in stability in *LARP1* KO HeLa cells. **(E)** Comparison of TOP score with 3’UTR length. Each point represents a gene. (A,B) boxes show 25^th^, 50^th^ and 75^th^ percentile, whiskers represent the largest/smallest values or 1.5 X the interquartile range. (A,B) TOP RNAs compared to “not target” group using Wilcoxon rank sum test with Benjamini-Hochberg p-value adjustment for multiple testing. (C,D,E) Correlation assessed by Kendall’s rank τ coefficient, trendline plotted using locally estimated scatterplot smoothing.

### TOP RNAs with short 3’UTRs and higher TOP scores depend on MTORC1 for their translation

MTORC1 pathway activation downstream of TCR stimulation promotes T cell growth and proliferation and is essential for T cell responses (Waickman and Powell 2012). MTORC1 promotes the translation of TOP RNAs by phosphorylation of LARP1 which inhibits its association with the RNA cap, and by phosphorylation of the translation inhibitor EIF4E binding protein 1 (4E-BP1) which then releases EIF4E allowing translation initiation (Tcherkezian et al. 2014; Fonseca et al. 2015; Hong et al. 2017; Jia et al. 2021; Hochstoeger et al. 2024). The phosphorylation status of LARP1 had not been examined in T cells, therefore, we investigated the MTORC1-dependent regulation of LARP1 in CD4^+^ T cells using proteomics and phospho-proteomics datasets from Tan *et al* (Tan et al. 2017a) who compared control and *Raptor* (Regulatory-Associated Protein of mTOR) KO naïve and activated CD4^+^ T cells. RAPTOR is an essential component of MTORC1 where it acts as a scaffold protein recruiting substrates for phosphorylation by the MTOR kinase. TCR stimulation induced LARP1 phosphorylation at two MTORC1 dependent sites: S743 and T476/T747 (Fig.8A). These phosphorylation sites regulate the interaction between LARP1 and RAPTOR (Jia et al. 2021). Thus we can confirm MTORC1 phosphorylation of LARP1 in activated T cells, but did not detect the phosphorylation sites associated with changes in RNA cap affinity.

**Figure 8.**
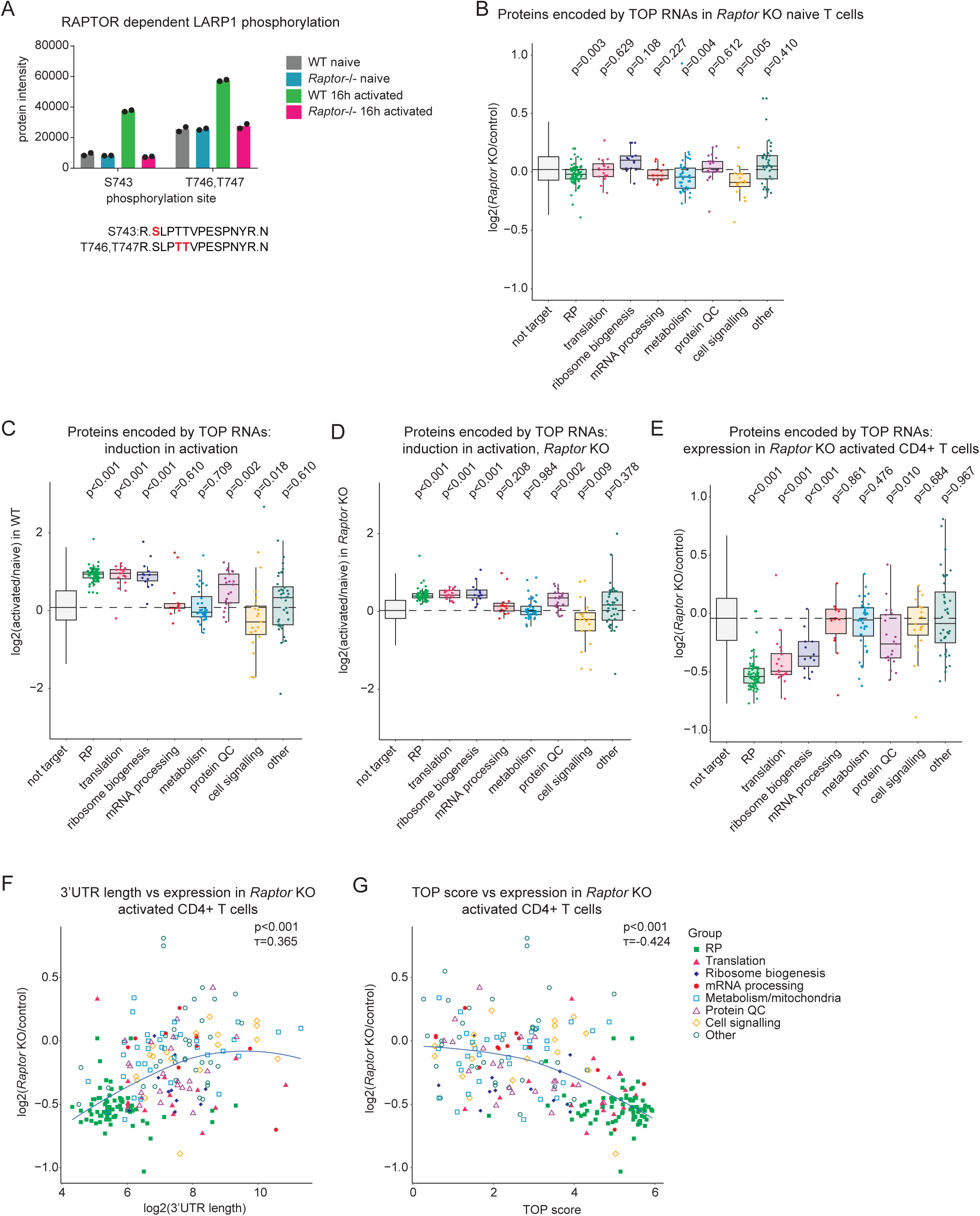
TOP RNAs with short 3’UTRs depend on the MTOR pathway for their translation. **(A)** RAPTOR dependent LARP1 phosphorylation induced by TCR and coreceptor stimulation determined by phosphoproteomics of control and *Raptor* KO naïve and activated mouse CD4^+^ T cells(Tan et al. 2017a). Bars represent the mean and points indicate biological replicates (n=2 mouse samples). Sequence context of the phosphorylation sites shown. **(B-E)** Differential expression of proteins encoded by TOP RNAs in naïve *Raptor* KO vs control CD4^+^ T cells (B), 16 hour activated vs naïve CD4^+^ T cells (C), 16 hour activated vs naïve *Raptor* KO CD4^+^ T cells (D), 16 hour activated *Raptor* KO vs control CD4^+^ T cells (E), each point represents a gene, boxes show 25^th^, 50^th^ and 75^th^ percentile, whiskers represent the largest/smallest values or 1.5 X the interquartile range from the box. Proteins encoded by TOP RNAs compared to “not target” group using Wilcoxon rank sum test with Benjamini-Hochberg p-value adjustment for multiple testing. **(F-G)** Comparison of 3’UTR length determined by Nanopore in activated CD4^+^ T cells (F) or TOP score (average terminal pyrimidine length) determined from FANTOM5 naïve CD4 cell CAGE data (G) with differential expression of proteins encoded by TOP RNAs in 16 hour activated control vs *Raptor* KO CD4^+^ T cells. Each point represents a gene, correlation assessed by Kendall’s rank τ coefficient, trendline plotted using locally estimated scatterplot smoothing.

In naive T cells, where MTORC1 signalling is low and LARP1 is not phosphorylated at the sites promoting interaction with RAPTOR, loss of *Raptor* had a subtle impact on the proteome overall (Fig.8B). Of the proteins encoded by TOP RNAs; the RPs and proteins involved in metabolism and cell signalling tended to be RAPTOR dependent, but the effect size was small (Fig.8B). Because it is activated downstream of TCR signalling, RAPTOR has a more prominent role in activated T cells. T cell activation increases the expression of many proteins encoded by TOP RNAs, including RPs, translation factors and proteins involved in ribosome biogenesis and protein quality control (Fig.8C). Notably not all TOP-encoded proteins are induced following activation and those involved in cell signalling instead tended to be repressed. In *Raptor* KO CD4^+^ T cells, the same groups of TOP-encoded proteins showed increased expression, but with a smaller fold change (Fig.8D). Thus, MTORC1 selectively promotes the expression of the TOP-encoded RPs, translation factors and proteins involved in ribosome biogenesis and protein quality control (Fig.8E). Since we found that TOP transcripts with short 3’UTRs and higher TOP scores were the most translationally repressed (Fig.5E, Fig.6C) we investigated the relationship between these measures and protein expression in *Raptor* KO activated T cells. Both short 3’UTR length and high TOP scores correlated with regulation via MTORC1, (Fig.8F-G, Fig.S1F-G). Thus, features at both the 3’ and 5’ end of TOP RNAs may link the differences we observed in TOP RNA translation and stability to altered regulation upon T cell activation.

## Discussion

T cell activation, triggered by T cell receptor (TCR) ligation, induces cell growth, proliferation and differentiation into effector T cells. Induction of ribosome biogenesis is crucial to increase the biosynthetic capacity of T cells during activation (Asmal et al. 2003; Allison et al. 2016; Tan et al. 2017b; Galloway et al. 2021; Rosenlehner et al. 2024). Ribosomal proteins are encoded by TOP RNAs, which are regulated by the RNA binding protein LARP1. RP transcripts are characterised by high stability and translational repression in naive T cells, followed by a MTORC1-dependent increase in translation following T cell activation. By mapping LARP1-RNA interactions in naïve and activated T cells using CLIP, we identified TOP RNAs with a broad range of functions including novel and T cell-specific transcripts. Interestingly, TOP RNAs with different functions had different patterns of regulation. Of the proteins encoded by TOP RNAs only those involved in ribosome biogenesis and protein translation, folding and turnover had increased expression following T cell activation. Proteins encoded by TOP RNAs that have functions in mRNA processing, metabolism and cell signalling were not induced and were not dependent on MTORC1 for their expression in activated T cells.

Among the TOP RNAs, RP transcripts had the lowest translation efficiencies, highest stabilities and greatest dependence on MTORC1 for their translation following T cell activation. We identified two key features of the RP transcripts that set them apart from other TOP RNAs: the first was their very short 3’UTRs which were typically between 30-60 nucleotides long. Shorter 3’UTRs were associated with both increased dependency on LARP1 for transcript stability, and MTORC1-dependent translational control. RPs also had the highest TOP scores; since this is a measure of average pyrimidine sequence length at the transcript start site it indicates that they have both a strong TOP sequence and high tendency for transcription to begin within it. The TOP score was associated with MTORC1-dependent translational control, but does not directly link to stability. These transcript sequence features may influence TOP RNA regulation to allow T cells to selectively induce the expression of specific groups of TOP RNA-encoded proteins following activation, whilst others were maintained at a stable level, or even repressed.

### LARP1 interactions during T cell activation

LARP1 has multiple RNA-interacting domains, and can also interact with the polyA tail indirectly through PABPC1. Most direct LARP1-mRNA interactions identified by CLIP in CD4^+^ T cells were interactions with TOP motifs at the 5’ end of transcripts, these are likely to involve the DM15 domain and La-module which directly recognise these elements. Loss of the ^m7^G RNA cap methyltransferase RNMT, its activating co-factor RAM, or first nucleotide O-2 methyltransferase CMTR1 leads to reduced TOP RNA expression in T cells suggesting an important role for the cap binding DM15 domain of LARP1 in TOP RNA stabilisation (Galloway et al. 2021; Knop et al. 2024; Galloway et al. 2025). The most repressed TOP RNAs in *Rnmt, Ram* and *Cmtr1* cKO CD4^+^ T cells are those encoding ribosomal proteins, which we found are the most dependent on LARP1 for transcript stabilisation. Although RPs had consistently high TOP scores, other TOPs tended to have lower scores. In some cases this was due to transcription start site heterogeneity resulting in a mixture of transcripts with the pyrimidine rich motif at the start, or further down in the 5’UTR; and in some cases it was due to interruption of the TOP sequence with a purine nucleotide. Higher TOP scores correlated with greater LARP1 binding indicating that the position or length of the pyrimidine stretch promotes the LARP1 interaction. Among transcripts with variable start sites, it is unclear whether LARP1 can bind all isoforms: the LARP1 LA-module can recognise uncapped oligopyrimidine sequences which could allow binding downstream of the TSS (Al-Ashtal et al. 2021). Additionally, mRNA starting with as few as two pyrimidines can be translationally regulated by LARP1 albeit to a lesser extent than those with longer pyrimidine stretches(Philippe et al. 2020). Together this suggests there is a degree of flexibility in TOP recognition.

Our data indicates that although LARP1 is predominantly associated with 40S ribosomal subunits there is still a substantial amount of polysome associated LARP1 in activated CD4^+^ T cells. Since the direct interaction between LARP1 and the 40S ribosome would block translation (Saba et al. 2024) LARP1 is most likely to be interacting with polysomal mRNA, either directly or through PABPC1. Indeed, mutation of the PABPC1 interacting domain of LARP1 reduces its interaction with mRNA as well as its association with polysomes in HeLa cells (Smith et al. 2021). Notably as well as mRNA that directly binds LARP1 through TOP motifs, there are non-TOP RNAs that appear in LARP1 RIP experiments that could interact solely through PABPC1 (Mura et al. 2015; Smith et al. 2021). The DM15 domain of LARP1 blocks EIF4F assembly to have an inhibitory effect on TOP RNA translation(Fonseca et al. 2015), so it was surprising that the binding of LARP1 close to the cap of TOP RNAs encoding RPs was equivalent in naïve and activated T cells, when the translation of RP transcripts is induced following activation. This could either reflect ongoing translational inhibition, binding via the LA module which would not occlude the cap, or a dynamic interaction of the DM15 domain that still allows for ribosome loading.

### Functions for LARP1 outside of translational efficiency and stability

A major question arising from our data is the function of the interaction of LARP1 with TOP RNAs with longer 3’UTRs and less stringent TOP scores. Is it simply to promote stability and regulate translation, or are there additional, context-dependent, functions to LARP1 binding? Interestingly, LARP1 has a role in localised translation at the mitochondrial outer membrane, and promotes oxidative phosphorylation, healthy cristae structure and mitochondrial quality control (Zhang et al. 2019; Ma et al. 2024; Bai et al. 2025). These observations were made across a range of organisms including fruit flies, nematodes and human cells, suggesting a well conserved function. Notably, the TOP RNAs encoding metabolic enzymes and mitochondrial proteins were among those with longer 3’UTRs and more moderate TOP scores that were not dependent on MTORC1 for their translation in activated T cells.

Recently LARP1 has also been shown to promote the synaptic enrichment of TOP RNAs, particularly RP transcripts, in neurons (Williams et al. 2025). This supports a broader role for LARP1 in localised translation. LARP1 is also reported to interact with components of the cytoskeleton (Burrows et al. 2010). We identified TOP RNAs encoding cytoskeletal proteins: Actg1 (Actin Gamma 1), Myl6 (Myosin light polypeptide 6), Tmsb4x (Thymosin beta-4) and Tmsb10 (Thymosin beta-10). Localised translation of these proteins could explain the impact of LARP1 on cytoskeletal morphology and cell migration (Burrows et al. 2010). If LARP1 does enable targeted translation of mRNAs to different subcellular locations, there must be additional factors involved to provide more specificity such as RNA binding proteins or proteins assembling on nascent peptides.

### Why is 3’UTR length associated with TOP RNA translation and stability?

There was little LARP1 binding to the 3’UTR regions, instead, clues to the molecular basis of the association between 3’UTR length with transcript stability could lie with the interaction between LARP1 and PABPC1. Transcripts with longer 3’UTRs are more prone to exon junction complex (EJC)-independent nonsense mediated decay (NMD), which is inhibited by PABPC1 (Munoz et al. 2023). LARP1 could inhibit EJC-independent NMD by stabilising the association between PABPC1 and the polyA tail either through direct binding or by increasing polyA tail length (Mattijssen et al. 2021; Ogami et al. 2022). This mechanism may be more effective on transcripts with shorter 3’UTRs that place PABPC1 closer to the terminating ribosome (Kajjo et al. 2022). The proximity between terminating ribosomes and PABPC1:LARP1 complex could also influence translation efficiency. In the closed loop model, interactions between EIF4G and PABPC1 bring the 3’ and 5’ ends of mRNA together, facilitating 40S ribosome recycling for subsequent rounds of translation (Thompson and Gilbert 2017; Vicens et al. 2018). Since LARP1 interacts with the 5’ and 3’ ends of mRNAs it has been portrayed in closed loop models, but usually in the context of translation repression (Berman et al. 2021). Because LARP1 also binds the 40S ribosome at the mRNA channel it could have a more direct role in recycling 40S subunits, pausing their re-recruitment into translation initiation complexes at the 5’ end of TOP RNAs (Saba et al. 2024; Wolin et al. 2025). Other RNA binding proteins are involved in translational repression of RP RNAs under conditions of MTORC1 inhibition including EIF4A1(Shichino et al. 2024) and EIF4E-BP (Hochstoeger et al. 2024; Wolin et al. 2025), thus the regulation of TOP translation may require the concerted action of multiple RNA binding proteins.

## Conclusions

We determined that LARP1 is induced following T cell activation, with the majority associating with 40S ribosomal subunits. Despite changes in translation of many TOP RNAs, LARP1-mRNA interactions are not dramatically remodelled on the majority of TOP RNAs following TCR stimulation. We identified TOP RNAs with diverse functions and different degrees of translational regulation and stabilisation. Short 3’UTR length and greater TOP scores were associated with MTORC1-sensitive translational repression and greater stabilisation by LARP1. mRNA encoding RPs are the prototypical TOP transcripts and were the most extreme examples, having the shortest 3’UTRs, highest TOP scores, highest stabilities, lowest translation efficiencies and greatest dependency on MTORC1 for protein expression following T cell activation. For non-RP TOP transcripts there are possibilities for additional roles for LARP1 beyond the regulation of translation and mRNA stability.

## Materials and Methods

### Mice

Mouse experiments were conducted in the Biological Resource Unit at the University of Dundee. Research using animals at the University of Dundee was reviewed by the Welfare and Ethical Use of Animals Committee, prior to authority being granted under the UK Home Office Animals (Scientific Procedures) Act 1986. All work was conducted under project licence PCF18BE491. Mice were housed in single sex groups with littermates where possible, with constant access to food (RM3, Special Diet Services) and water, on a 12:12 h light:dark schedule, at 21°C. Mice were used when they were between 8 and 14 weeks old and were cre^-^ control mice from our *Rnmt*^fl/fl^ colony (Galloway et al. 2021) which is maintained on a C57Bl/6J background. Both female and male mice were used as indicated for individual experiments, sample size calculation, blinding and randomisation were not carried out for our study design. Mice were culled by exposure to a rising concentration of CO_2_ and death confirmed by severing the femoral artery.

### Preparation of CD4^+^ T cells

Lymph nodes (inguinal, brachial, axillary, superficial cervical, mesenteric, lumbar, caudal), were dissected from mice, and mashed through a 70µm cell strainer (Falcon) to prepare cell suspensions. CD4^+^ T cells were magnet sorted using the EasySep mouse CD4^+^ T cell isolation kit (Stemcell Technologies, catalog #19852). Cells were counted and purity was assessed by incubating cells with FC block (anti mouse CD16/32 clone 93, Biolegend), anti-CD4 FITC (clone RM4-5, Biolegend), anti-Thy1.2 APC (clone 53–2.1, Biolegend), and 0.1 µg/ml DAPI in FACS buffer (PBS +2% FCS (Gibco) then analysed using the BD FACSVerse (BD Biosciences) or Novocyte (Acea Biosciences) flow cytometer. To generate activated CD4^+^ T cells, tissue culture treated plates were coated with anti-mouse CD3ε antibody (clone 145-2C11, Biolegend, 5µg/ml) and anti-mouse CD28 antibody (clone 37.51, Biolegend, 1µg/ml) in PBS for two hours at 37°C or overnight at 4°C. Plates were rinsed with PBS then CD4^+^ T cells were added at 1million/ml in T cell culture medium (RPMI + 10% heat inactivated FCS + pen/strep + 50µM 2ME).

### Western blotting

1million CD4^+^ T cells from 12 week old female mice were lysed directly in Laemmli buffer (50mM Tris pH 6.8, 2% SDS, 10% glycerol, 100mM DTT, bromophenol blue). Proteins were resolved by SDS-PAGE on an 8% polyacrylamide gel then transferred onto PVDF membranes (Millipore) with Tris-glycine buffer (25mM Tris, 190mM glycine, 20% methanol). Membranes were incubated with anti-Actin rabbit monoclonal antibody (clone EPR16769, Abcam) or anti-LARP1 rabbit polyclonal antibody (Protein Tech, 13708-1-AP), followed by the HRP-conjugated Goat anti-Rabbit IgG (H+L) Secondary Antibody (Thermo Fisher Scientific, 31460) and developed with Pierce Super signal ECL (Thermo Fisher Scientific), then imaged using an ImageQuant LAS 4000 (GE healthcare). Western blots were quantified using NIH Image J software (v1.53).

### Ribo Mega-SEC

Ribo Mega-SEC was carried out as part of our *Rnmt* cKO study and the data are available at PXD023832(Galloway et al. 2021). Briefly, 3.9x 10^6^ 20 hour activated CD4^+^ T cells from 9 week old male mice (three replicate samples each using two mice) were lysed in polysome extraction buffer (20 mM Hepes-NaOH (pH 7.4), 130 mM NaCl, 10 mM MgCl_2_, 5% glycerol, 1% CHAPS, 0.2 mg/ml heparin, 2.5 mM DTT, 20 U SUPERase In RNase inhibitor, cOmplete EDTA-free Protease inhibitor), filtered, then passed through a SEC column (Agilent Bio SEC-5, 2,000 Å pore size, 7.8 × 300 mm with 5 μm particles) using a Dionex Ultimate 3,000 Bio-RS uHPLC system (Thermo Fisher Scientific). UV absorbance was monitored at 260nm. Proteins from each fraction were treated with Benzonase, reduced using TCEP (25 mM final concentration), and alkylated using N-Ethylmaleimide (25 mM final concentration). Proteins were purified using SP3 hydrophilic and hydrophobic beads, digested with trypsin, re-purified with the SP3 beads then resuspended in 50 μl of 1% formic acid then analysed using a Q-exactive plus (Thermo Fisher Scientific) mass spectrometer. Further details are available in our original publication(Galloway et al. 2021).

Data from the three control mouse biological replicates, consisting of heavy polysome, light polysome/80S, 60S, 40S and “small 1” and “small 2” fractions were analysed. Raw data were analysed in Maxquant using the mouse Swissprot database for peptide identification. iBAQ scores for each protein were used for further analysis in R. Data from each fraction and replicate were analysed using the sum of iBAQ scores as a normaliser between replicates. Boxplot graphs were drawn using ggplot2.

### CLIP

CLIP was carried out as described in the seCLIP protocol with a few modifications (Van Nostrand et al. 2017; Galloway et al. 2021). Naïve and activated CD4^+^ T cells from 11 week old male mice were suspended in PBS and exposed to 400mJ of 254nm UV then snap frozen on dry ice. The naïve CD4^+^ T cell sample was made from four mice and the activated CD4^+^ T cell sample was a single mouse. Cells were thawed into lysis buffer (50 mM Tris-HCl pH 7.4, 100 mM NaCl, 1% NP-40 (Igepal CA630), 0.1% SDS, 0.5% sodium deoxycholate, 1:200 Protease Inhibitor Cocktail III (Sigma), 1:200 murine RNAse inhibitor (NEB)) and lysed on ice for 15 minutes. Lysates were incubated with 1µl/ml turbo DNAse (Thermo Fisher Scientific) and 1µl/ml of RNAse1 diluted 1:2000 in PBS (Thermo Fisher Scientific) for 5 mins at 37°C then returned to ice before centrifugation to remove insoluble material.

Lysates were precleared with protein G dynabeads (Thermo Fisher Scientific) at 4°C for 1hour. Lysates underwent immunoprecipitation with 50µl of protein G dynabeads preincubated with 4 µg of either anti-LARP1 rabbit polyclonal antibody (Abcam, ab86359), or Rabbit IgG isotype control (Thermo Fisher Scientific, 31235) overnight at 4°C, 10% of the LARP1 samples were taken for the size matched input control. Beads were washed three times for 5 minutes with high salt wash buffer (50 mM Tris-HCl pH 7.4, 1 M NaCl, 1 mM EDTA, 1% NP-40, 0.1% SDS, 0.5% sodium deoxycholate) with 2M urea. Samples were then washed with lysis buffer, then TAP buffer (10 mM Tris pH 7.5, 5 mM MgCl_2_,100 mM KCl, 0.02% Triton X-100). RNA was dephosphorylated with FastAP Thermosensitive Alkaline Phosphatase (Thermo Fisher Scientific) at 1200 rpm, 37°C, then beads washed in wash buffer (20 mM Tris-HCl pH 7.4, 10 mM MgCl_2_, 0.2% Tween-20, in water) twice. A 5% aliquot of the immunoprecipitate was labelled with gamma 32P-ATP (Perkin Elmer) using PNK (NEB) to check for RNA-protein complexes. For library construction, 2.5µl of the 3’ RNA adapter InvRiL19 (40µM) (sequences in Table S3) was ligated to bead bound RNA fragments at room temperature for 75 minutes using the following mixture: 9µl H_2_O, 3µl 10X ligase buffer (500 mM Tris-HCl pH 7.5, 100 mM _MgCl2)_, 0.3 μL 0.1 M ATP, 0.8µl 100% DMSO, 9µl 50% PEG 8000, 0.4µl Murine RNase Inhibitor, 2.5 μL High concentration T4 RNA Ligase (NEB). Beads were then washed in wash buffer, high salt wash buffer, then wash buffer.

RNA-protein complexes were denatured in NuPage loading buffer (Thermo Fisher Scientific) + 50mM DTT, resolved in 3-8% Tris Acetate gel (Thermo Fisher Scientific) in Tris Acetate SDS buffer (Thermo Fisher Scientific), and transferred to a nitrocellulose membrane in NuPage transfer buffer (Thermo Fisher Scientific). RNA protein complexes were identified from the 32P-ATP labelled membrane, and cut from the membrane containing libraries. RNA was released by incubating with Proteinase K for 20 minutes at 37°C, then with 420mg/ml urea in Proteinase K buffer for 20 minutes at 37°C and purified by phenol chloroform extraction.

The input library was dephosphorylated with FastAP Thermosensitive Alkaline Phosphatase (Thermo Fisher Scientific) at 1200 rpm, 37°C, then cleaned up with 20µl myOneSilane beads (Thermo Fisher Scientific) using RLT buffer (Qiagen), NaCl, and ethanol in the binding buffer, and 75% ethanol as a wash buffer then dissolved made up in 10µl water. For 3’ adapter ligation 5µl of the input was mixed 0.5 μL InvRiL19 and 1.5µl DMSO then incubated for 75 minutes at room temperature with the following mixture: 1.5µl H_2_O, 2µl 10X ligase buffer, 0.2 μL 0.1 M ATP, 0.3µl 100% DMSO, 8µl 50% PEG 8000, 0.2µl Murine RNase Inhibitor, 1.3 μL High concentration T4 RNA Ligase (NEB). Input samples were then cleaned up again with 20µl myOneSilane beads, with RLT buffer plus ethanol as the binding buffer and 75% ethanol as the wash buffer.

RNA was reverse transcribed using superscript III (Thermo Fisher Scientific), excess primers removed with EXOSAPIT (Thermo Fisher Scientific), and RNA removed by heating at 70°C for 12 minutes with 120mM NAOH, followed by neutralisation with HCl. cDNA was cleaned up with 10µl myOneSilane beads using RLT buffer (Qiagen) with ethanol as a binding buffer and eluted in 5 μL 5 mM Tris-HCl pH 7.5. 0.8µl of the 3’ DNA linker InvRand3Tr3 and 1µl DMSO were added then ligated at room temperature overnight with this mixture: 1.5µl High concentration T4 ligase (NEB), 2µl proprietary T4 ligase buffer, 0.2µl 0.1M ATP, 9µl of 50% PEG 8000, and 1.1µl H_2_O.

Adapter linked cDNA was cleaned up with silane beads, and amplified by PCR with Phusion High-Fidelity DNA Polymerase (Thermo Fisher Scientific) using the HF buffer provided. PCR products were cleaned up with AMPure beads, and gel purified on an agarose gel, extracted with a Qiagen MinElute gel extraction kit. Libraries were quantified using the qbit HS DNA assay (Thermo Fisher Scientific) and quality checked using the tapestation HS DNA assay (Agilent).

Libraries were sequenced with a Nextseq High Output v2.5 kit (75 cycles, single end).

### CLIP analysis

Naïve and activated CD4^+^ T cell LARP1 CLIP data were analysed alongside a second naïve CD4^+^ T cell dataset from our previous publication(Galloway et al. 2021) available at NCBI GEO with accession GSE160325. The second naïve CD4^+^ T cell dataset is from four female mice. These libraries were prepared and sequenced together. 10nt randomer barcodes were extracted using umi_tools, then adapter sequences were trimmed from the reads using cutadapt. Reads were aligned to ribosomal RNA (rRNA) from a customised Fa file available from GSE128947: Supplementary_Data_1.fa using bowtie2(Langmead and Salzberg 2012). Reads which did not align to the rRNA were aligned to the mm39 mouse genome with gencode vM33 basic annotation using STAR 2.7.11(Dobin et al. 2013). PCR duplicates were removed using umicollapse(Liu 2019). Aligned CLIP data were analysed using htseq-clip (Sahadevan et al. 2023). Annotation was converted to the bed format using --splitExons option to divide exons into 5’UTR, 3’UTR and CDS. Sliding windows were created using the standard window size of 50 nucleotides and step size 20 nucleotides. Count tables were analysed using DEWseq in R (Schwarzl et al. 2024). Windows with a significant enrichment in the naïve CD4^+^ T cell LARP1 CLIP datasets over their respective size matched controls were identified using the local fit type and bionomial Likelihood ratio test options with an adjusted p value threshold of 0.01. False positives were identified by comparing the IgG CLIP and input, with and adjusted p value threshold of 0.05. False positive windows were removed from the list of hits. For each gene the window with the lowest p value was selected and represented in Fig.2A, sequences were extracted from these windows and analysed using MEME(Bailey et al. 2015). The LARP1 binding profile was generated using Deeptools (Ramirez et al. 2016): bamCoverage was used with -- Offset 1 1 so that the position of the read start was considered, the matrix was generated using the metagene option to join exons and profile plotted with plotProfile. Reads aligning to individual genes and the 18S rRNA were plotted using Gviz in R. To compare the CLIP read density to transcriptome read density in Fig.4, control naïve (Galloway et al. 2021) and activated (Knop et al. 2024) T cell dataset from our previous studies available at NCBI GEO with accessions GSE160267 and GSE198541 were used. Naïve and 20 hour activated CD4^+^ T cell data were aligned to mm10 with gencode vM12 or vM25 basic annotation respectively, using STAR version 2.5.2b(Dobin et al. 2013). Read count tables generated by STAR were normalised to transcript length using the longest transcript length per gene, and used to calculate transcripts per million. TOP transcripts were defined as those with LARP1 binding sites in the 5’UTR and assigned to functional groups for further analysis. For the STRING analysis (Szklarczyk et al. 2023), connections represent the “physical subnetwork” i.e. physical interactions between the proteins has been reported.

### Ribosome footprinting analysis

Naïve T cell ribosome footprinting data(Khalsa et al. 2019) from NCBI GEO dataset series GSE128298 was downloaded using sra_tools and activated CD4^+^ T cell ribosome footprinting data (Galloway et al. 2021) is available at NCBI GEO with accession GSE160326. Adapter sequences were trimmed out using cutadapt, and sequences aligning to ribosomal RNA removed using bowtie 2 (Langmead and Salzberg 2012). The data were then aligned to the mm39 mouse genome with gencode vM33 basic annotation using STAR 2.7.11 (Dobin et al. 2013). Reads aligning to coding sequence (CDS) regions were counted using HTseq (Putri et al. 2022). Counts per million for each gene were calculated in R and TE calculated as CPM ribosome footprints / CPM total RNA. The naïve T cell data consists of two replicates containing a mix of CD4^+^ and CD8^+^ T cells, the activated CD4^+^ T cell data consists of three replicates each from two female mice. For each dataset the mean translation efficiency across the replicates was calculated.

### Nanopore sequencing analysis

Nanopore sequencing of activated CD4^+^ T cells was published previously(Galloway et al. 2025) and is available at NCBI GEO with accession GSE284393, four control genotype datasets were used, each replicate is from a single mouse, two are females and two are males. Full length cDNA sequences were aligned to the mm10 mouse genome using flair align, splice junctions corrected using flair correct with gencode vM25 annotation, isoforms collapsed using flair collapse with the stringent option, then quantified using flair-quantify(Tang et al. 2020). To determine the 3’UTR lengths of the most common isoform of each gene, a mean average count per million for the four replicates was calculated in R, and the transcript variant with the highest CPM selected, only transcripts with a CPM greater than 5 were considered. 3’UTR length was measured from the transcript end to nearest stop codon. Images showing transcript variants were exported from IGV (version 1.14.1).

### TOP score analysis

TOP scores were calculated using the C57Bl/6J mouse naive CD4 T cell HeliScopeCAGE data (n=1) from the FANTOM5 project aligned to mm10 (Abugessaisa et al. 2017; Noguchi et al. 2017), (Lizio et al. 2015). The scoring was carried out as described by Phillipe et al (Philippe et al. 2020) using the code associated with their paper (https://github.com/carsonthoreen/tss_tools) to analyse the first 6 nucleotides from each transcription start site. A filter of at least 50 CAGE reads was applied and where more than one transcript variant was identified for a given gene, the transcript variant with the highest TOP score was selected.

### RNA stability and proteomics analysis

Naïve CD4^+^ T cell transcript stabilities were taken from supplemental table 3 from Hwang et al 2020(Hwang et al. 2020). These transcript stabilities were measured by actinomycin D pulse chase.

*LARP1* KO vs control HeLa cell transcript stabilities were taken from supplemental table 4 from Hochstoeger et al 2024(Hochstoeger et al. 2024). These transcript stabilities are the average of two independent clones of the LARP1 WT and LARP1 KO cell lines and were measured by 4-thiouridine labelling.

*Raptor*^-/-^ and control naïve and activated CD4^+^ T cell proteomics data was taken from Tan et al 2017(Tan et al. 2017a). The *Raptor*^-/-^ and control naïve and activated CD4^+^ T cell data were taken from supplemental table 5A and the average changes between the two biological mouse replicates are used. The expression of LARP1 at 0, 2, 8 and 16 hours was taken from supplemental table 1A and the LARP1 phospho proteomics data was taken from supplemental table 5B.

### Graphs and statistical analysis

Statistical comparisons were calculated in R (v4.5.2) using the Kendall rank correlation test or Wilcoxon rank sum test with Benjamini-Hochberg p-value adjustment. Graphs were plotted in R or GraphPad Prism 7.

## Acknowledgements

We thank the Biological Resource Unit, Flow Cytometry Facility, Fingerprints Proteomics Facility and Tayside Centre for Genomic Analysis at the University of Dundee, and core services and computational biology department at the CRUK Scotland Institute. The manuscript was critically reviewed by Catherine Winchester (CRUK Scotland Institute), Axel Arthur (CRUK Scotland Institute), Martin Turner (Babraham Institute) and Louise Matheson (Babraham Institute).

## Data availability

CLIP data generated in this study is available on the NCBI GEO repository with accession number: GSE198483. No new code or reagents were generated for this study.

## Author contributions

Conceptualization, A.G. and V.H.C.; investigation, A.G.; writing, A.G. and V.H.C; supervision, V.H.C.

## Figure Legends

**Figure S1. Trends in TOP RNA attributes without ribosomal protein transcripts**

Protein-coding transcripts with LARP1 binding sites in the 5’UTR were divided into groups depending on their function, in these analyses TOP RNAs encoding ribosomal proteins were excluded.

**(A)** Comparison of 3’UTR length by Nanopore in activated CD4^+^ T cells with naïve T cell translation efficiency.

**(B)** Comparison of TOP score (average 5’ terminal pyrimidine length) of LARP1-bound RNAs determined from FANTOM5 naïve CD4^+^ T cell CAGE data with naïve T cell translation efficiency.

**(C)** Comparison of 3’UTR length with change in transcript stability in *LARP1* KO HeLa cells.

**(D)** Comparison of TOP score with change in transcript stability in *LARP1* KO HeLa cells.

**(E)** Comparison of 3’UTR length with TOP score.

**(F)** Comparison of 3’UTR length with differential expression of proteins encoded by TOP RNAs in 16 hour activated control vs *Raptor* KO CD4^+^ T cells.

**(G)** Comparison of TOP score with differential expression of proteins encoded by TOP RNAs in 16 hour activated control vs *Raptor* KO CD4^+^ T cells.

TE (translation efficiency: ribosome protected fragment RNA counts per million (CPM)/total mRNA CPM). Each point represents a gene, correlation assessed by Kendall’s rank τ coefficient, trendline plotted using locally estimated scatterplot smoothing.

